# Crohn’s patients and healthy infants share immunodominant B cell response to commensal flagellin peptide epitopes

**DOI:** 10.1101/2023.08.08.552496

**Authors:** Qing Zhao, Lennard Wayne Duck, John T. Killian, Alexander F. Rosenberg, Peter J. Mannon, R. Glenn King, Lee A. Denson, Subra Kugathasan, Edward N. Janoff, Maria C. Jenmalm, Charles O. Elson

**Author notes:** Corresponding author. Qing Zhao Charles O. Elson.

## Abstract

About half of patients with Crohn’s disease (CD) develop selective serum IgG response to flagellin proteins of the *Lachnospiraceae* family. Here, we identified a dominant B cell peptide epitope in CD, locating in the highly conserved “hinge region” between the D0 and D1 domains at the amino-terminus of *Lachnospiraceae* flagellins. Serum IgG reactive to this epitope is present at an elevated level in adult CD patients and in pediatric CD patients at diagnosis. Most importantly, high levels of serum IgG to the hinge epitope were found in most infants from 3 different geographic regions (Uganda, Sweden, and the USA) at one year of age. This vigorous homeostatic response decrements with age as it is not present in healthy adults. These data identify a distinct subset of CD patients, united by a shared reactivity to this dominant flagellin epitope that may represent failure of a homeostatic response beginning in infancy.

## INTRODUCTION

Inflammatory bowel disease (IBD) is characterized as chronic inflammation in the gut mucosa, with Crohn’s disease (CD) and ulcerative colitis (UC) as the two main forms. It is generally accepted that aberrant immune responses to environmental factors, especially to the gut microbiota, play a critical role in the pathogenesis of IBD(*1–3*). Although CD and UC share various genetic polymorphisms(*4*), dominant antigens identified in CD are considerably different from that in UC(*5–7*). Flagellins expressed by commensal *Lachnospiraceae* bacteria and CD4^+^ T cells specific to these flagellins are found in blood and intestinal lesions of Crohn’s patients(*8, 9*), suggesting that *Lachnospiraceae* flagellin is a keystone antigen driving gut inflammation in at least a subset of Crohn’s patients. Serum IgG reactivity to CBir1, a *Lachnospiraceae* flagellin isolated from mice, is elevated and associated with disease complications in patients with CD, but not in patients with UC or healthy controls (HC)(*6, 8*). In addition, anti-flagellin IgG reactivity was shown to predict the development of CD years prior to disease onset(*10–13*). Therefore, sero-reactivity to *Lachnospiraceae* flagellins has been used as a biomarker for CD diagnosis. However, although *Lachnospiraceae* flagellin has been identified as an immunodominant antigen in CD at the protein level, the dominant B cell peptide epitopes and where they are located remain unclear.

The motility of various commensal and pathogenic bacteria, especially those that belong to the Proteobacteria and Firmicutes (reclassified as Bacillota now) phyla, is driven by their surface expression of flagella, which can consist of as many as 30,000 flagellin monomers(*14*). Although flagellin proteins have a vast span of diversity among different organisms, they share a characteristic molecular structure consisting of amino- and carboxyl-domains (D0 and D1) that are conserved within a given bacterial family(*15, 16*). The amino- and carboxyl-domains are separated by a hypervariable region (D2 and D3 domains). In addition to conferring motility, flagellin is also one of the most potent immune activators known in the microbiota(*17–21*). As a result, flagellin proteins expressed by intestinal pathogens such as *Salmonella* spp. and *Escherichia coli* (*E. coli*) serve as major constituents mediating intestinal infection and inflammation(*22, 23*). However, immune reactivity targeting flagellins expressed by commensal bacteria, especially those belong to the *Lachnospiraceae* family, is only elevated in patients with CD, but not in patients with UC or adult HC, despite the presence of abundant intestinal *Lachnospiraceae* bacteria in the latter(*24*).

Previously, using a microbiota protein antigen microarray, we have shown that ∼30% of CD patients in our regional IBD cohort recruited at the University of Alabama at Birmingham (UAB) had elevated serum IgG reactivity to more than 10 different flagellins expressed by *Lachnospiraceae* bacteria (from both human and mouse origin), and reactivity to individual flagellins was highly correlated to each other(*24*), suggesting that serum IgG response to *Lachnospiraceae* flagellins was driven by reactivity to shared epitopes, which remained unidentified. Consistently, these multi-flagellin reactive patients had significantly increased flagellin-specific effector/memory CD4^+^ T cells in the circulation(*24*), which could travel back to the gut and drive colitis (*25–27*). We proposed that patients with multi-flagellin reactivity are more likely to develop disease complications in CD, and early identification of these patients through reactivity to the shared epitopes would greatly help with their disease treatment.

Here, using a novel flagellin peptide microarray and a flagellin peptide cytometric bead array assay, we report the identification of a key IgG B cell epitope in *Lachnospiraceae* flagellins in CD patients, which locates at the “hinge region” between the D0 and D1 domain in the highly conserved amino-terminus (N-term) of *Lachnospiraceae* flagellins. Sero-reactivity to the dominant epitope was identified in our regional adult CD patients, and in a multi-site-recruited cohort of pediatric CD patients. In addition, the magnitude of this epitope-specific reactivity was positively correlated with multi-flagellin reactivity in CD patients, and predicted the development of a complicated disease course in pediatric CD patients 2-3 years in advance. Interestingly, we observed a robust level of serum IgG response to this dominant epitope in over 90% of infants of diverse geographic origins, which peaked at one year of age, indicating that the elevated IgG reactivity to *Lachnospiraceae* flagellins in CD patients may result from an aberrant immune recall response of sensitization that occurred in early life.

## RESULTS

### Cohort description

Three cohorts were used in this study, including a discovery cohort of adult IBD patients and HC recruited at UAB(*24*), a validation cohort of treatment-naïve pediatric CD patients and non-IBD controls collected at 28 sites in the USA and Canada (the RISK Stratification Study)(*28*), and a healthy and non-IBD control cohort of infants and their mothers recruited in Uganda, the USA(*29*), and Sweden(*30*).

### *Lachnospiraceae* flagellin peptide microarray identifies key IgG binding epitopes in CD patients

With our previous microbiota protein microarray, Pearson correlation analysis showed that serum IgG response to most *Lachnospiraceae* flagellins strongly correlated with each other in CD patients(*24*). This observation suggested that serum IgG reactivity in CD patients was targeting shared epitopes among different *Lachnospiraceae* flagellins. To test this hypothesis, we generated a novel flagellin peptide microarray consisting of 1512 peptides derived from 19 different *Lachnospiraceae* flagellins of both human and mouse origin. These were sequential 15mer peptides overlapping by 5 amino acids, which covered the entire sequences of 19 flagellin proteins. Sera from CD patients, HC, and UC patients were probed against the array, and peptide-specific IgG or IgA reactivities were detected with fluorescently labeled secondary antibodies (**Figure 1A**). In the heatmap where serum IgG reactivity to peptides in the D0 and D1 domains of the N-term of *Lachnospiraceae* flagellins is shown (**Figure 1B**), those in CD patients clustered apart from those in UC patients and HC, whereas the response to *Lachnospiraceae* flagellin peptides in UC and HC subjects was not different. Notably, reactivities from CD sera fell in two major clusters, with increased IgG binding identified at p26-41 and p41-61 between the D0 and D1 domains in the amino chain of *Lachnospiraceae* flagellins (**Figure 1B and 1C**), which was not seen in UC or HC subjects. There was no significant difference seen in serum IgG reactivity among CD, UC, and HC subjects against other peptides included in the microarray (**Supplementary** Figure 1). Differences in the IgA response to *Lachnospiraceae* flagellin peptides among groups were not observed. Sequence analysis and motif discovery analysis revealed that, regardless of human or mouse origin, the majority of the 19 *Lachnospiraceae* flagellins included in the peptide microarray have significant amino acid homology in both the N-term and C-term of their D0 and D1 domains. Specifically, such homology peaks at the “hinge region” between the D0 and D1 domains at the N-term (**Figure 1D and 1E**), which is significantly different from flagellins expressed by Proteobacteria *Salmonella dublin* (*S. dublin*) and *E. coli* (**Supplementary Table 1**), although the latter two are homologous to each other. Multiple likely B cell epitopes were predicted in this hinge region when *Bepipred Linear Epitope Prediction 2.0* was performed through Immune Epitope Database(*31*), consistent with this region being highly immunogenic (**Supplementary Table 2**). The dominant B cell epitope in *Lachnospiraceae* flagellins identified in CD patients with the flagellin peptide microarray locates right at this “hinge region”. It includes sub-epitopes p25-44, which locates at the D0 domain, and p41-59 which locates at the D1 domain. Hereafter, we will refer to the dominant *Lachnospiraceae* flagellin B cell epitope in CD, D0-1N p25-59, as the “hinge peptide”.

**Figure 1.**
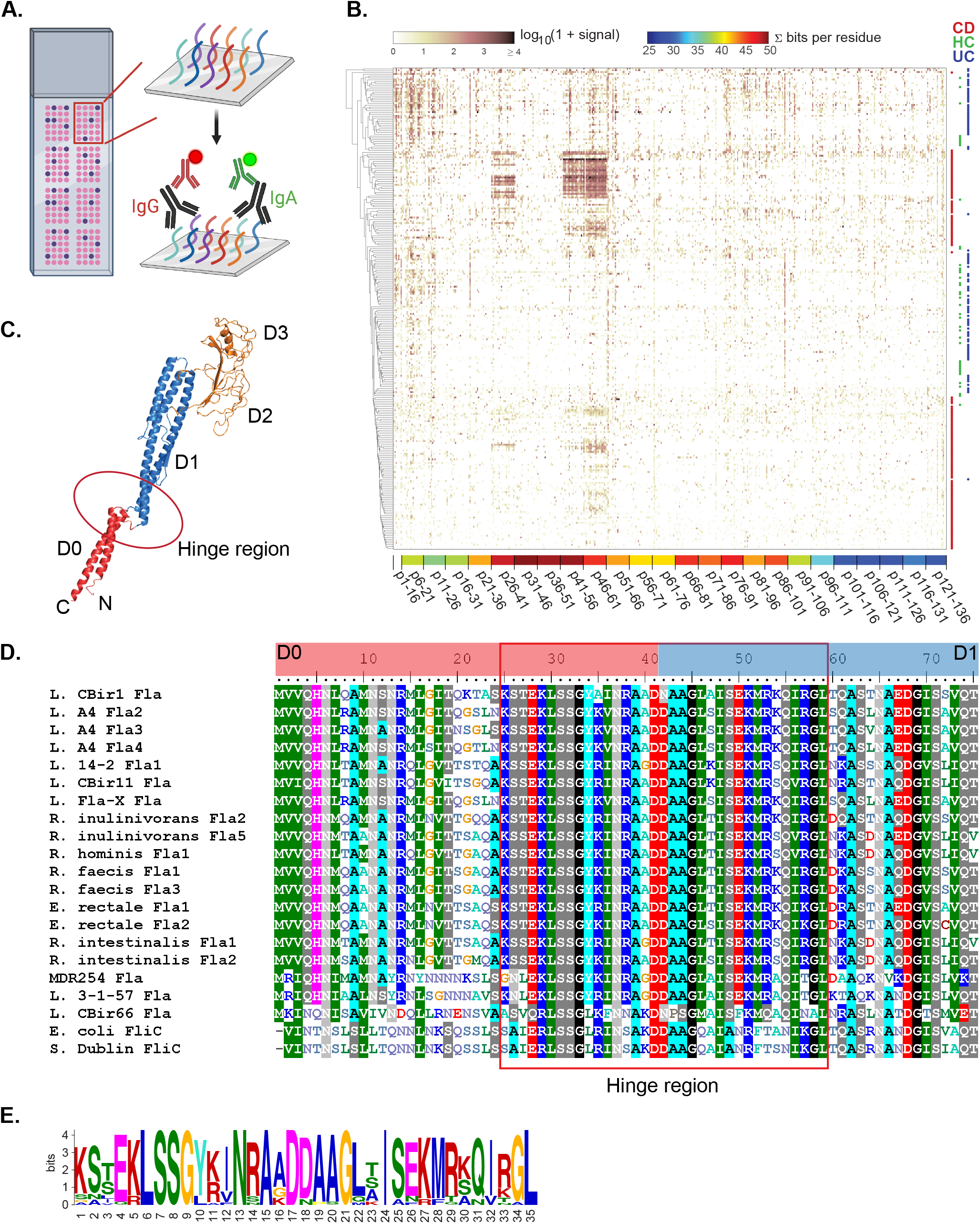
*Lachnospiraceae* flagellin peptide microarray identifies key IgG binding epitopes in CD patients. **(A)** Schematic strategy of the flagellin peptide microarray. **(B)** Heatmap showing serum IgG reactivity for sequential, overlapping fifteen-residue flagellin peptides (108 peptides from each of the 19 different flagellins) in CD (n=135), HC (n=42), and UC (n=88) subjects. Sera were diluted at 1:100. Rows (subjects) were clustered using Euclidean distance and complete linkage applied to log-transformed data. Columns (peptides) are sorted by peptide position (and include peptides from up to 19 different flagellins per position). Cohort groups are indicated by colored dots on the right margin. Peptide starting position (relative to the N-terminus) is indicated on the bottom. For each 15-mer segment, per-residue information content is computed at each position across all flagellins (except *L. CBir11* Fla and *L. CBir66* Fla) and summated to indicate sequence conservation, shown as color. Although all peptides were used for the clustering, only peptides up to and including 121-136 are depicted in this heatmap. (See **Supplementary** Figure 1 for version showing all peptides). **(C)** 3D structure of *L. A4* Fla2 as a representative flagellin showing the location of the hinge region. **(D)** Sequence alignment of flagellins included in the peptide microarray plus *S. dublin* FliC and *E. coli* FliC. Sequences were aligned in BioEdit using the ClustalW alignment feature. Flagellin D0 and D1 domains in the N-term are indicated by the color bars on top, whereas the hinge region is marked by the red box. **(E)** Motif discovery of the flagellin hinge region.

### Serum IgG multi-flagellin reactivity in CD patients is driven by reactivity to the dominant *Lachnospiraceae* flagellin peptide epitope

In order to confirm the findings of the flagellin peptide microarray, we developed a flagellin peptide cytometric bead array, where serum IgG, IgA, and IgM reactivity to different bead-bound antigens can be simultaneously detected in a multiplexed fashion (**Figure 2A**). Eight biotinylated conserved peptides (D0N p1-19, D0N p25-44, D1N p41-59, D0-1N p25-59, D1N p74-102, D1C p391-407, D0-1C p410-435, D0C p437-460) covering the N-term and C-term of D0 and D1 domains of *Lachnospiraceae* flagellins were synthesized and individually conjugated to 4-micron fluorescent beads (**Figure 2B and 2C**). Beads conjugated with different peptides can be differentiated by their fluorescent intensity in the APC/Cy7 channel with flow cytometry (**Figure 2C**). Serum samples (UAB cohort) from CD patients with multi-flagellin reactivity on the microbiota protein microarray (≥10 different flagellins, CDhigh), CD patients who reacted to less than 10 or no *Lachnospiraceae* flagellins (CDlow), UC patients and HC were assayed and IgG, IgA, and IgM reactivity to individual peptides were detected with fluorescently labeled secondary antibodies. Standard curves using human IgA, IgM, and IgG capture beads and known concentrations of polyclonal human IgA, IgM, and IgG were obtained and applied to quantify peptide-specific antibodies, allowing cross-comparison of samples run at different times. Data from this cytometric bead array assay strongly validated the results we obtained from the flagellin peptide microarray (**Figure 2D**). IgG reactivity to the “hinge peptide” (D0-1N p25-59) is highly represented in CDhigh patients compared to CDlow, UC, and HC (**Figure 2D-F**). In addition, CDhigh patients have significantly elevated serum IgG specific to the sub-epitopes (D0N p25-44 and D1N p41-59) compared to CDlow patients, UC patients, and HC (**Figure 2G and 2H**), but not to other conserved peptides of the *Lachnospiraceae* flagellins including D0N p1-19, D1N p74-102, D1C p391-407, and D0-1C p410-435 (**Figure 2I-K**, **and 2M**). These data indicate that multi-flagellin IgG reactivity in Crohn’s patients is driven by reactivity to the hinge region of the N-terminus. Of note, CDhigh patients also have significantly elevated serum IgG reactivity to D0C p437-460 (**Figure 2L**), which is the terminal conserved region of *Lachnospiraceae* flagellin C-terminus, although the magnitude of reactivity is much lower than that to the N-term hinge peptide. Serum IgA and IgM responses to individual conserved peptides were also analyzed, but consistent with the microbiota protein microarray and flagellin peptide microarray data, no differential reactivity among groups was observed.

**Figure 2.**
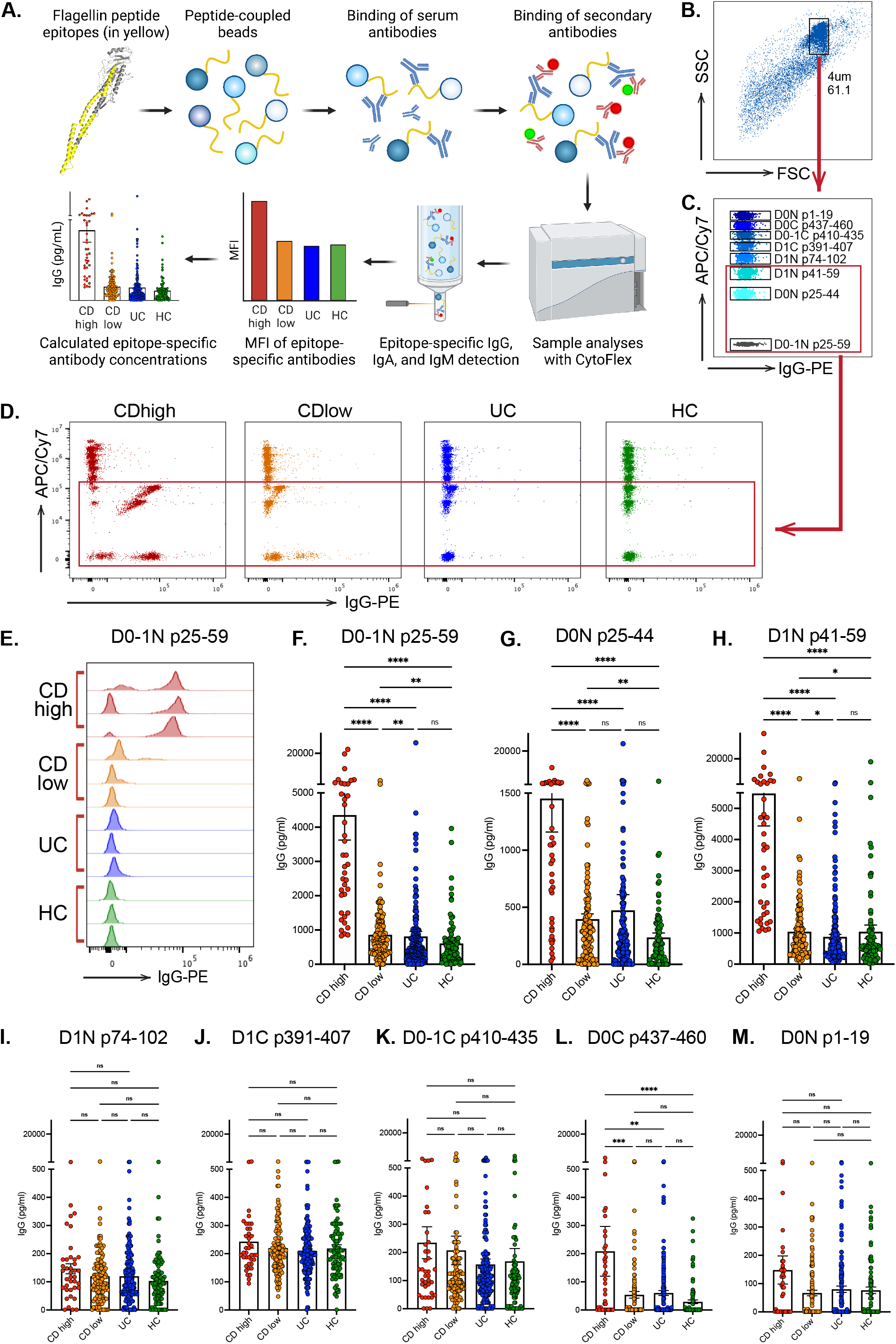
Serum IgG multi-flagellin reactivity in CD patients is driven by reactivity to the dominant *Lachnospiraceae* flagellin peptide epitope. **(A)** Schematic strategy of the flagellin peptide cytometric bead array. **(B and C)** Gating strategies of the flagellin peptide cytometric bead array. **(D)** Sera from CDhigh (n=39), CDlow (n=126), UC (n=181), and HC (n=95) subjects in the UAB cohort were probed against the flagellin peptide cytometric bead array. Representative flow plots of serum IgG reactivity to flagellin peptide epitopes in each group are shown. The red box highlights the hinge region peptides. **(E)** Histogram overlay of serum IgG reactivity to *Lachnospiraceae* D0-1N p25-59 in 3 representative CDhigh, CDlow, UC, and HC, respectively. **(F-M)** Epitope-specific IgG concentration in the serum of CDhigh, CDlow, UC, and HC subjects to indicated flagellin peptide epitopes. Data are presented as means ± SEM and analyzed with nonparametric Kruskal-Wallis test. *P<0.05, **P<0.01, ***P<0.001, ****P<0.0001.

### *Lachnospiraceae* hinge peptide is the dominant B cell epitope in pediatric CD patients and predicts the development of disease complications

Although we were able to confidently identify the dominant B cell peptide epitope in CD patients using our IBD cohort recruited at UAB, there were several limitations involved. First, it is a regional adult cohort of IBD, which might not represent a broader IBD population. Also, the refractory nature of this cohort (rate of biologic therapy in CD was 78% and rate of surgery in CD was 66%)(*24*) limited the power to analyze the association between the epitope-specific IgG sero-response and disease complications in CD. Therefore, we decided to validate this finding using a multi-site-recruited pediatric cohort consisting of patients who were newly diagnosed with CD as well as non-IBD controls (the RISK Stratification Study)(*28*). CD patients in this cohort represented the variety of disease phenotypes regarding disease location, severity, and behavior, and they were recruited at diagnosis prior to any treatment. Furthermore, patients were followed up to 36 months after the initial sample collection, and their disease course was well-documented, thus the prospective nature of this cohort enabled us to test whether serum IgG to the dominant epitope in CD patients was able to predict the development of disease complications later. In specific, we referred to the Montreal classification and defined developing stricturing (B2) or penetrating (B3) disease behavior as a complicated disease course.

To confirm that pediatric CD patients in the RISK cohort are reactive to *Lachnospiraceae* flagellins, we first performed the microbiota protein microarray as previously described(*24*). Sera at recruitment of CD patients who remained inflammatory (B1 behavior) at 3-year follow-up, CD patients who developed strictures or penetrating disease (B2/3 behavior) at 3-year follow-up, and non-IBD controls were probed with 19 *Lachnospiraceae* flagellins of human and mouse origin, in addition to *S. dublin* FliC and *E. coli* FliC. Previous study using this cohort has reported an augmented serum IgG specific to *L. CBir1* Fla in 35% of CD patients who remained B1 at follow-up, and in 62% of CD patients who developed B2/3 at follow-up(*28*). Consistent with this data, our results showed that CD patients that remained B1 exhibited a trend of increased, although not significant, serum IgG reactivity to *Lachnospiraceae* flagellins when compared to control subjects (**Figure 3A**). In contrast, CD patients that developed B2/3 at follow-up showed significantly elevated serum IgG reactivity to the majority of *Lachnospiraceae* flagellins compared to controls and to patients who remained B1 (**Figure 3A and Supplementary** Figure 2). Consistent with what we observed in our adult CD cohort recruited at UAB, a subset of pediatric CD patients also exhibited serum IgG reactivity to the majority of *Lachnospiraceae* flagellins tested (**Figure 3B**), and the proportion of patients that were multi-flagellin reactive was significantly higher in those who developed B2/3 at follow-up (**Figure 3C and 3D**).

**Figure 3.**
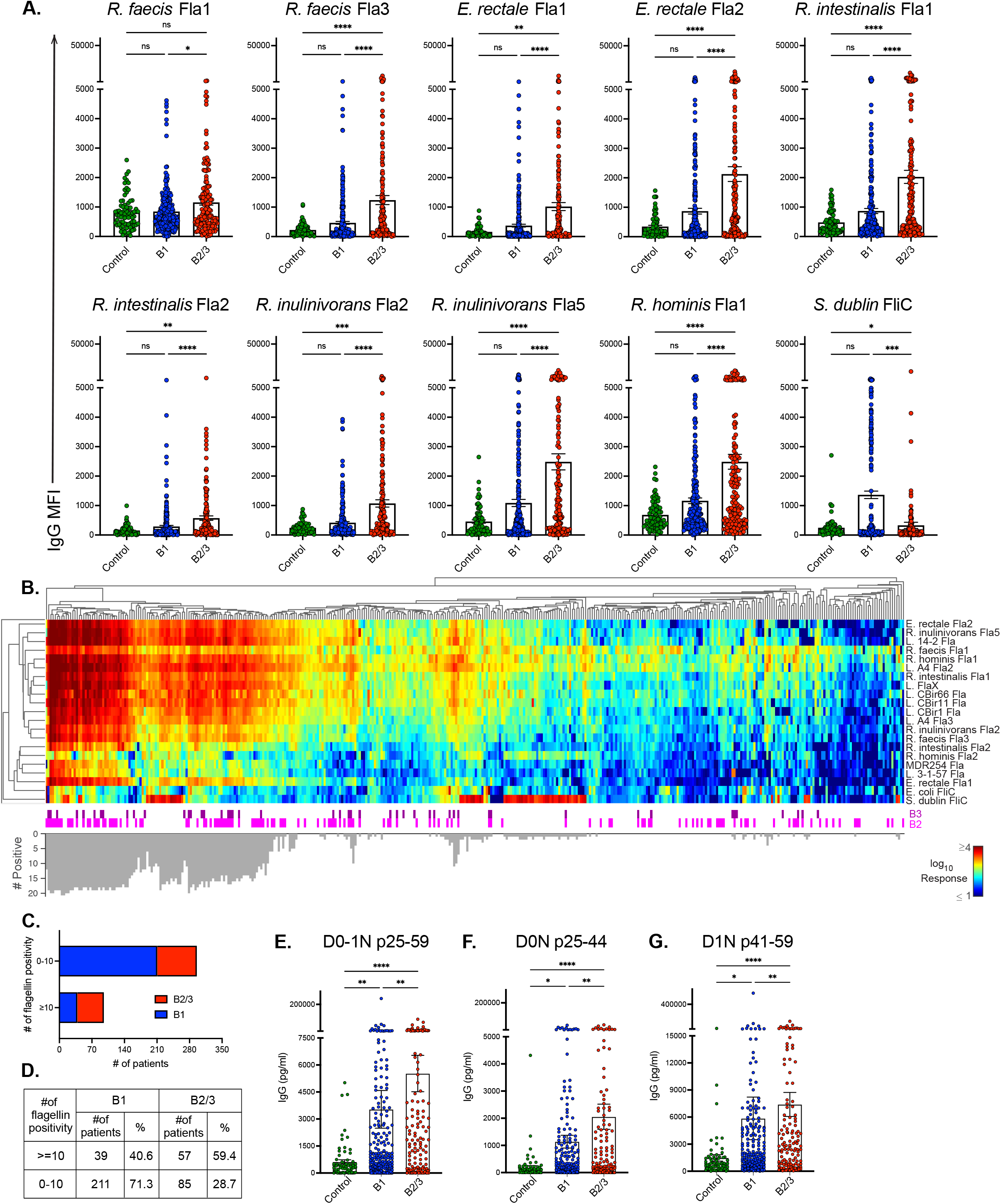
*Lachnospiraceae* hinge peptide is the dominant B cell epitope in pediatric CD patients and predicts the development of disease complications. **(A)** Sera of CD patients (grouped by disease behavior at 3-year follow-up using the Montreal classification, n=250 for B1 (inflammatory), n=142 for B2 (stricturing) or B3 (penetrating)) and non-IBD controls (n=72) from the RISK cohort were probed against the microbiota protein microarray. Mean fluorescent intensity (MFI) of serum IgG specific for indicated flagellin antigens is presented as means ± SEM and analyzed with nonparametric Kruskal-Wallis test. *P<0.05, **P<0.01, ***P<0.001, ****P<0.0001. **(B)** Heatmap of log_10_ IgG response (color) on the microbiota protein microarray for 21 flagellins (rows) and 392 individuals with CD in the RISK cohort (columns). Rows and columns were hierarchically clustered using Euclidean distance on the log data, average linkage and optimal leaf ordering maximizing the sum of similarity between every leaf and all other leaves in the adjacent cluster. Montreal disease classification at follow-up (B2 and B3) is indicated below the heatmap, with unmarked individuals being B1. The bar graph at the bottom indicates the number of positive flagellin responses for each individual, where a positive response is two standard deviations above the mean (non-log transformed) of a non-IBD control cohort (n=72, not shown) per flagellin. **(C and D)** Summary of multi-flagellin reactivity in CD patients with B1 or B2/3 behavior at follow-up. **(E-G)** Sera of CD patients (grouped by disease behavior at 3-year follow-up, n=249 for B1, n=140 for B2/3) and non-IBD controls (n=72) from the RISK cohort were probed against the flagellin peptide cytometric bead array. Epitope-specific IgG concentration in different groups to indicated flagellin peptide epitopes is shown. Data are presented as means ± SEM and analyzed with nonparametric Kruskal-Wallis test. *P<0.05, **P<0.01, ****P<0.0001.

Next, we tested whether the *Lachnospiraceae* flagellin hinge peptide also served as the dominant B cell epitope in CD patients in the RISK cohort by performing the flagellin peptide microarray with sera of the multi-flagellin reactive CD patients. Similar to what we observed in our adult IBD cohort recruited at UAB, serum IgG response in multi-flagellin reactive CD patients in the RISK cohort mainly targeted p26-41 and p41-61 between the D0 and D1 domains in the N-term of *Lachnospiraceae* flagellins (**Supplementary** Figure 3). Finally, patient and control sera from the whole cohort were assayed with the flagellin peptide cytometric bead array. Similar to adult CD patients, the dominant serum IgG binding sites in pediatric CD patients located at the conserved *Lachnospiraceae* flagellin hinge region between D0 and D1 domains in the N-term (D0-1N p25-59, D0N p25-44, and D1N p41-59, **Figure 3E-G**). IgG sero-reactivity to other peptides was either not significantly different among groups (D1N p74-102 and D1C p391-407, **Supplementary** Figure 4) or at a much lesser magnitude compared to the hinge peptide (D0-1C p410-435, D0C p437-460, and D0N p1-19, **Supplementary** Figure 4). Moreover, response to the N-term hinge peptide in patients who developed B2/3 at follow-up was significantly higher than that of patients who remained B1 at follow-up, indicating the prognostic value of hinge peptide specific serum IgG in CD patients early on, even at initial disease diagnosis. In conclusion, the dominant B cell peptide epitope in *Lachnospiraceae* flagellins in CD patients were validated with both the flagellin peptide microarray and flagellin peptide cytometric bead array, using the RISK cohort which represents a broad variety of disease phenotypes and behaviors. More importantly, results from the flagellin peptide cytometric bead array demonstrated that elevated serum IgG to *Lachnospiraceae* flagellin hinge peptide was associated with stricturing or penetrating disease at 3-year follow-up, thus can be used to predict the development of disease complications in CD.

### Serum IgG to *Lachnospiraceae* hinge peptide strongly correlates with multi-flagellin reactivity in CD

Next, we sought to determine whether serum IgG reactivity to the conserved peptides included in the flagellin peptide cytometric bead array correlates with reactivities to *Lachnospiraceae* flagellins included in the microbiota protein microarray. First, we performed Pearson correlation analysis between serum IgG reactivity against individual flagellins and serum IgG reactivity specific to individual peptide epitopes for CD, HC, and UC subjects from the UAB cohort and CD patients from the RISK cohort (**Figure 4A**). The most significant correlations with serum reactivities specific to *Lachnospiraceae* flagellin proteins were observed in serum reactivities against D0N p25-44, D1N p41-59, and D0-1N p25-59, i.e. the hinge peptide, especially in CD patients from the UAB cohort and the RISK cohort. Sero-reactivities to the hinge peptide did not universally correlate with all flagellins tested, in that the correlation with that against Proteobacteria flagellins such as *E. coli* FliC and *S. dublin* FliC was significantly reduced. Moreover, sero-reactivity to the C-term conserved epitopes D0-1C p410-435 and D0C p437-460 showed slight significance in correlating with that to *Lachnospiraceae* flagellin proteins in RISK cohort CD patients. To better view the correlation in each subject, we plotted individual sero-reactivity against the most significant peptide, D0-1N p25-59, and one of the least significant peptides, D1C p391-407, with that of *R. intestinalis* Fla1, and *E. coli* FliC, respectively (**Figure 4B**). Reactivity to D0-1N p25-59, especially in CD patients from both UAB and the RISK cohort correlated significantly with that of *R. intestinalis* Fla1, which was expressed by bacteria in the *Lachnospiraceae* family, but poorly with *E. coli* FliC. In contrast, reactivity to D1C p391-407 did not correlate with that of *R. intestinalis* Fla1 in any subject group, demonstrating that reactivity to the hinge peptide, but not to other conserved flagellin peptides, underlies sero-reactivity to *Lachnospiraceae* flagellins. Furthermore, a good concordance was found between sero-reactivity to the hinge peptide obtained from the flagellin peptide cytometric bead array and flagellin sero-positivity determined by the microbiota protein microarray in both the UAB cohort and the RISK cohort (**Figure 4C and 4D**). This further demonstrates that multi-flagellin reactivity in CD patients is largely due to IgG binding to the dominant hinge peptide epitope.

**Figure 4.**
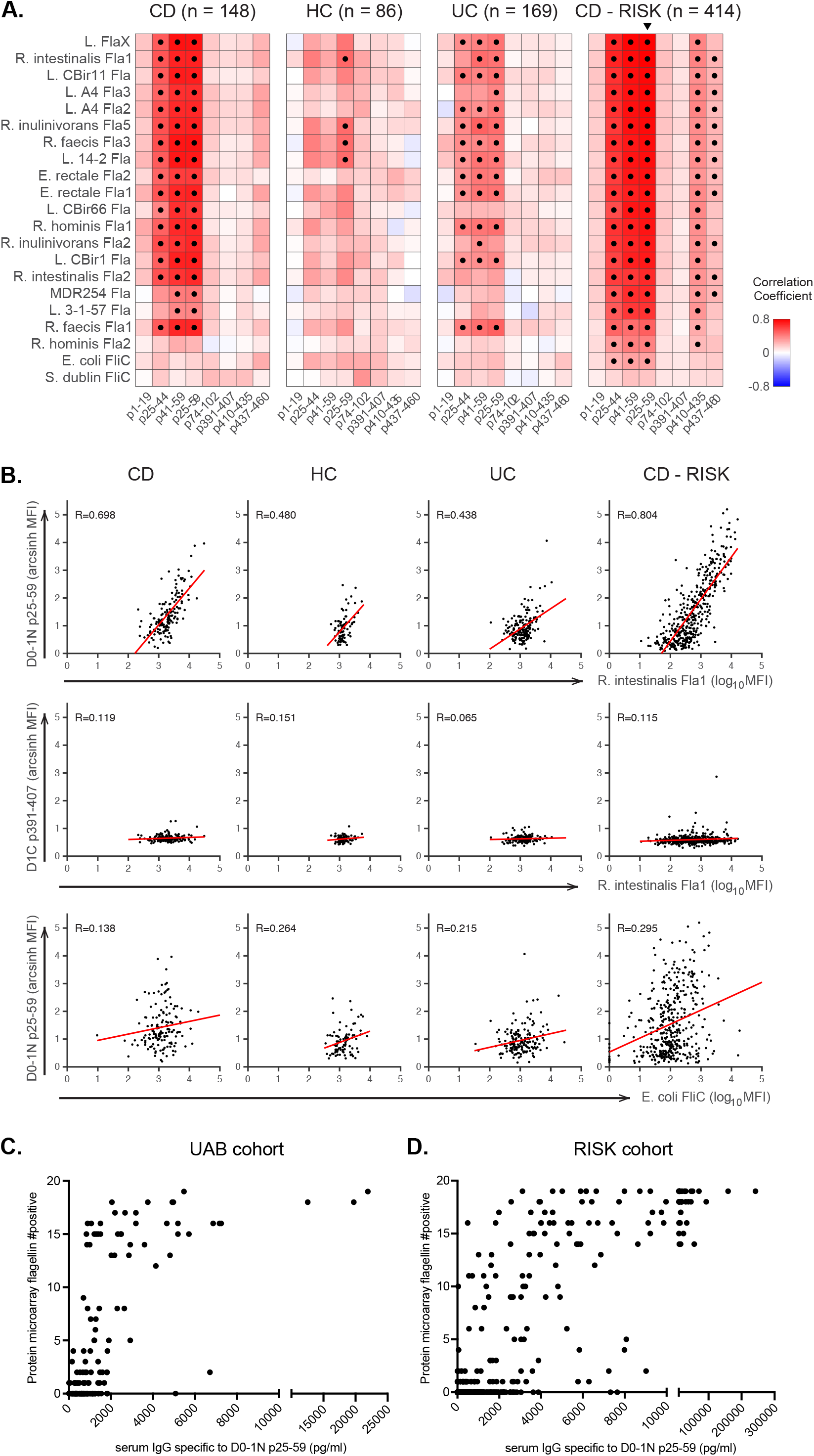
Serum IgG to *Lachnospiraceae* hinge peptide strongly correlates with multi-flagellin reactivity in CD. **(A)** Heatmaps representing correlation coefficients between serum IgG anti-flagellin reactivity (log_10_ MFI from microbiota protein microarray; rows) and serum IgG anti-peptide reactivity (hyperbolic arcsin MFI from flagellin peptide cytometric bead array; columns). Four separate heatmaps are shown for CD (n=148), HC (n=86), and UC (n=169) participants from the UAB cohort and CD (n=414) participants from the RISK cohort, respectively. Color corresponds to Pearson correlation coefficient indicated in the bar scale. Dots indicate correlations that were below Bonferroni-corrected p-value of 0.01 (based on 21 flagellins, 8 peptides, 4 groups = 672 tests) testing the hypothesis of no correlation against the alternative that there is a non-zero correlation. Rows (flagellins) are sorted based on the correlation for peptide D0-1N p25-59 in CD patients in the RISK cohort, as indicated by the triangle. **(B)** Hyperbolic arcsin MFI of serum IgG anti-*Lachnospiraceae* D0-1N p25-59 and D1C p391-407 of individual CD, HC, and UC participants from the UAB cohort and CD participants from the RISK cohort obtained from the flagellin peptide cytometric bead array was plotted against reactivity (log_10_ MFI) to *R. intestinalis* Fla1 and *E. coli* FliC obtained from the microbiota protein microarray, respectively. Least squares regression was performed, with the correlation line in red and R value shown for each plot. **(C and D)** The value of hinge-specific IgG obtained from the flagellin peptide cytometric bead array is plotted on X axis, and the number of flagellins bound by sera in the same individual CD patients on the microbiota protein microarray on the Y axis. Data from the UAB cohort and the RISK cohort are presented as indicated.

### Serum IgG reactivity to *Lachnospiraceae* flagellin hinge peptide is a homeostatic response in most infants

Our previous study showed a surprisingly similar level of serum IgG reactivity to Firmicutes flagellins in healthy and allergic Swedish infants at 2 years of age compared to that in adult CD patients(*30*). We wondered whether this high level of flagellin reactivity existed universally in infants raised in distinct environments. To test this hypothesis, sera were obtained from HIV-exposed but uninfected healthy infants from Uganda, healthy and allergic infants from Sweden, and healthy infants from the USA at 6 and 12 months of age, as well as their cord blood representing antibody reactivity of the mothers. Data from the Swedish and the American infants were grouped together, representing reactivity from developed countries, whereas data from the Ugandan infants represented reactivity in developing countries.

Serum samples were probed with the flagellin peptide cytometric bead array and the level of IgG reactivity to the hinge peptide in each individual at different time points was linked and compared to that of CD patients in both the UAB cohort and RISK cohort. Hinge peptide specific IgG response in the cord blood of both Uganda and US/Sweden cohorts was low and comparable to that identified in adult HCs from the UAB cohort (**Figure 5A**). Consistent with our previous microbiota protein microarray data(*30*), serum IgG response to the hinge peptide was significantly elevated at 12 months of age in both cohorts (**Figure 5A**), indicating that IgG sero-reactivity to *Lachnospiraceae* flagellins is a homeostatic response present in infants regardless of environmental exposures, and this heightened response is driven by reactivity to the hinge epitope. Accompanied with this, there was an increase in sero-reactivity to other microbiota antigens at 12 months of age(*30*), reflecting the development of homeostatic IgG response to endogenous gut microbiota in early life. Interestingly, increase in hinge-specific sero-reactivity already happened at 6 months of age in Ugandan infants, but remained low in infants in the US/Sweden cohort; whereas reactivity was much greater in US/Sweden infants than Ugandan infants at 12 months of age (**Figure 5A**). Furthermore, sero-reactivity to the hinge peptide was attributed to both sub-epitopes (D0N p25-44 and D1N p41-59) in the US/Sweden cohort (**Figure 5B and 5C**), but there was a lack of potent sero-reactivity to sub-epitope D0N p25-44 in Uganda infants at 12 months of age (**Figure 5B**). Distinct from response to the hinge peptide, serum IgG reactivity to the other peptide epitopes tested in the cytometric bead array in Ugandan infants exhibited a significant increase at 6 months of age and started to decline at 12 months of age, which was not observed in the US/Sweden cohort (**Figure 5A-C and Supplementary** Figure 5). Collectively, these data indicate that there is an environmental contribution to the kinetics of the development of homeostatic IgG response to the gut microbiota.

**Figure 5.**
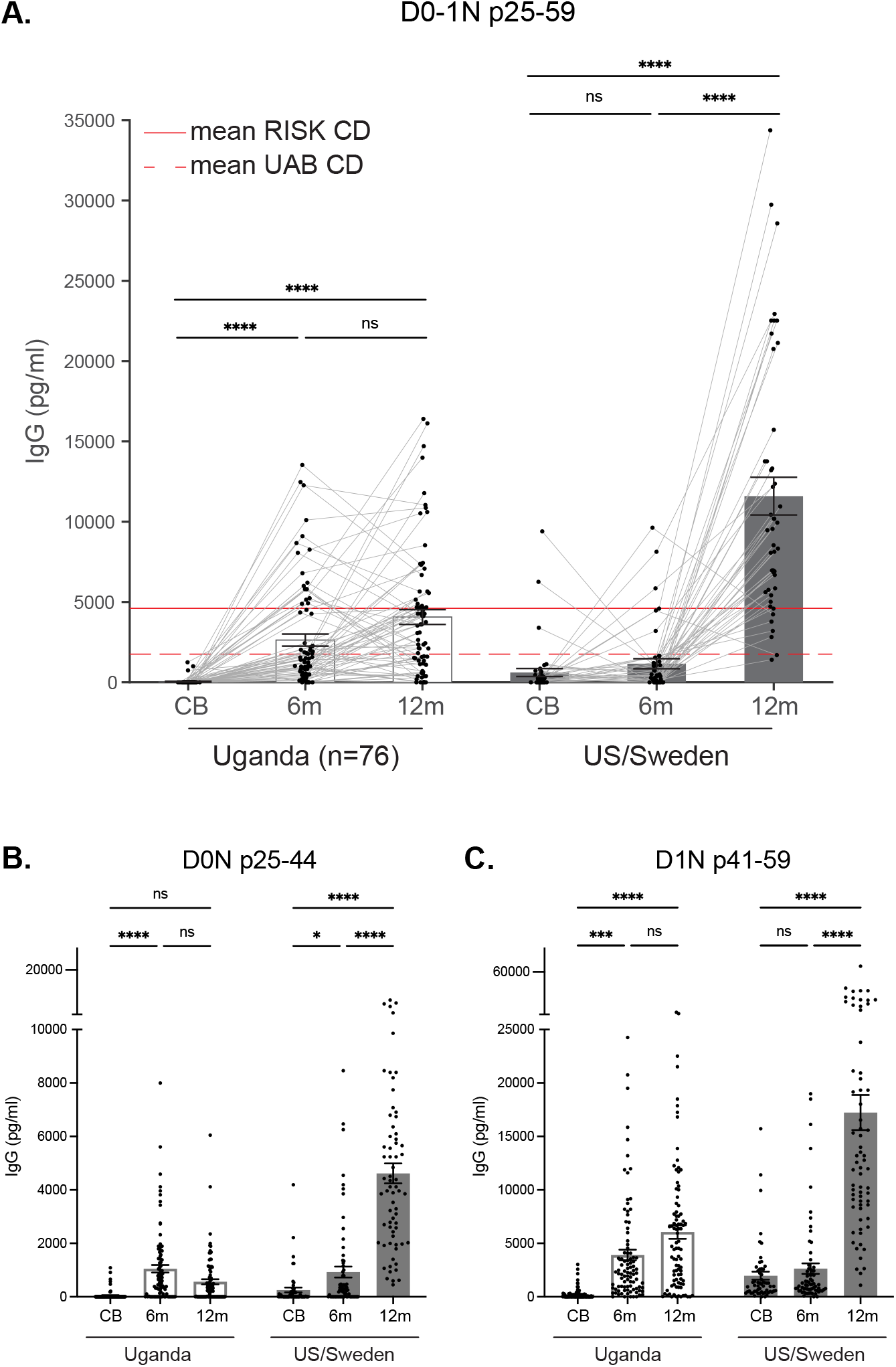
Serum IgG reactivity to *Lachnospiraceae* flagellin hinge peptide is a homeostatic response in most infants. **(A)** Sera of healthy and non-IBD control infants at 0 month (cord blood, CB), 6 months, and 12 months of age (with all three timepoints) from the Uganda cohort (n=76) and the US (n=10) /Sweden (n=36) cohort were probed against the flagellin peptide cytometric bead array. Concentration of serum IgG specific to *Lachnospiraceae* flagellin D0-1N p25-59 in different groups is shown. Each dot represents an individual and responses of the same individual at different timepoints are linked with the grey lines. The means of serum IgG response to *Lachnospiraceae* flagellin D0-1N p25-59 of CD patients in the UAB (broken line) and RISK (solid line) cohort are plotted. Data are presented as means ± SEM and analyzed with repeated measures ANOVA and Friedman test. ****P<0.0001. **(B and C)** Sera of healthy and non-IBD control infants at indicated timepoints (including individuals not represented in all three timepoints) from the Uganda cohort (n=92) and the US (n=16) /Sweden (n=54) cohort were probed against the flagellin peptide cytometric bead array. IgG concentration specific to *Lachnospiraceae* D0N p25-44 and D1N p41-59 sub-epitopes in different groups is shown, respectively. Data are presented as means ± SEM and analyzed with two-way ANOVA and Tukey’s multiple comparisons test. *P<0.05, ***P<0.001, ****P<0.0001.

## DISCUSSION

The dynamic interactions of the host with its microbiota begins at birth and continues throughout the life of the host. The composition of human gut microbiota changes over a lifetime, but most critically early in life where the microbiota is unstable until 3 years of age(*32–34*). A collection of genes is important in establishing and maintaining homeostasis with the microbiota, and when such genes are deleted from mice, colitis ensues(*35–37*). A similar process occurs in humans resulting in IBD, which eventuates in immune reactivity to the microbiota. Indeed, serologic responses to gut microbial antigens have been identified, such as perinuclear Anti-Neutrophil Cytoplasmic Antibodies (pANCA) in UC(*5, 38*), and anti-*Saccharomyces cerevisiae* antibodies (ASCA) in CD(*39, 40*). In addition, sero-response to flagellins, which is preferentially directed at flagellins of the *Lachnospiraceae* family of bacteria, was shown elevated specifically in patients with CD, but not in UC patients or HC(*6, 8, 24*). Data in this paper provides further insight into such response, identifying a shared dominant B cell peptide epitope in CD patients for the first time. Importantly, heightened sero-reactivity to this dominant epitope at CD diagnosis is able to predict the future development of disease complications up to 3 years, making the *Lachnospiraceae* flagellin hinge peptide a promising biomarker candidate for both CD diagnosis and prognosis.

IgG^+^ plasma cells only constitute a small proportion of plasma cells in the intestinal lamina propria at homeostasis, with most of the plasma cells being IgA^+^(*^41-43^*). However, anti-commensal IgG responses are significantly induced during intestinal inflammation, both in mouse models of colitis(*25, 44*), and in patients with IBD(*45–48*). It has been shown that IgG antibodies specific to flagellins derived from *Salmonella* spp. or *E. coli* aggravated colitis progression in mouse models(*48, 49*). However, these microorganisms are neither abundant mucus colonizers in healthy human intestines(*33, 43*), nor primary targets of serum IgG from CD patients(*24*). *Lachnospiraceae* of the Firmicutes phylum are one of the most abundant mucus-associated bacterial families in both human and mouse intestines(*33, 50*). They are very diverse: analysis of the genome of 273 isolates from human donors found only 60% of genes shared among multiple species(*51*). At homeostasis, they are beneficial commensals due to their ability to convert dietary fibers into short-chain fatty acids and thus promote the differentiation and maintenance of epithelial cells and regulatory CD4^+^ T cells (Treg) in the gut(*52, 53*). They are obligate anaerobes which are resistant to host antimicrobial peptides(*54*), and they produce lantibiotics that contribute to resistance to pathogen colonization, protecting the host(*51*). Many, but not all, are flagellated, a property facilitating their colonization of the mucosa. Therefore, flagellin, the monomeric subunit of the flagellum, is one of the most expressed bacterial proteins in the intestinal lumen and the mucosal layer. Crystallization studies of flagellin have revealed four domains, D0 and D1 that are localized internally in the flagellum, and D2 and D3 that are on the exposed surface of the flagellum(*16*). A recent study by Bourgonje *et al*. showed that serum IgG response to conserved motifs (D0 and D1 domains) in both the N-terminus and C-terminus of *Lachnospiraceae* flagellins was overrepresented in CD patients. Such response was identified using a massive antigen library including microbial peptides, immune epitope and food allergens using a high-throughput phage-display immunoprecipitation sequencing technique(*55*). Our study confirmed this finding, and beyond that, we defined the dominant B cell peptide epitope (D0-1N p25-59) of *Lachnospiraceae* flagellins in CD patients for the first time. Another interesting finding was that little or no reactivity to the Toll-like receptor 5 (TLR5) binding site of the flagellin (D1N p74-102) was detected in any of the subjects tested. As one of the receptors for flagellins, TLR5 is expressed by epithelial cells, including Paneth cells at the base of crypts, innate immune cells and some Treg cells(*56*). The TLR5 binding site on the flagellin locates in the D1 domain(*18, 57*), where only the monomeric form of flagellin is able to bind and activate host TLR5; those in the flagellum with D1 domain internalized thus do not bind TLR5. Antibodies targeting the conserved D0/D1 region were shown to neutralize flagellin TLR5 activation(*58*), but it is unclear which epitope contributed to the neutralization. It is possible that antibody directly blocking the flagellin TLR5 binding site inhibits its TLR5 activation. However, antibodies targeting the flagellin TLR5 allosteric binding site (located in the D0 domain)(*59–61*) may also alter its TLR5 activating function. Considering that the dominant B cell epitope identified in this study locates at the hinge region of D0-D1 domain of *Lachnospiraceae* flagellins, one can ask what role these antibodies may play in flagellin TLR5 activation in sero-positive subjects. In addition, it has been shown that the level of *Lachnospiraceae* bacteria was significantly reduced preceding the onset of CD and in newly diagnosed, treatment-naïve pediatric CD patients(*62, 63*), raising the question whether flagellin-specific antibodies in patients with CD are involved in the depletion of gut *Lachnospiraceae* bacteria. Do antibodies specific to D0N p25-44 and D1N p41-59 sub-epitopes play different roles? These are important questions that need to be further investigated in future studies.

Another important finding of this study was that most human infants at one year of age have a strong IgG response to *Lachnospiraceae* flagellin, targeting the exact dominant peptide epitope identified in CD patients. CD4^+^ T cell help is required for B cell responses to peptides, and T cells are restricted by major histocompatibility complex (MHC) class II molecules. The almost universal IgG response to the hinge peptide despite human immune diversity indicates that the T cell response is not restricted to any MHC class II allele. Indeed, T cell epitopes that are promiscuous, i.e., can bind to multiple MHC class II alleles, are enriched in *Lachnospiraceae* flagellins. Starting from 6 months, infants are often first introduced to solid food and thus experiencing a massive shift in the composition of the gut microbiota(*32, 33*). This is also a critical time window for the colonization and diversification of Firmicutes in the human gut. The increased serum IgG response to *Lachnospiraceae* flagellins reflects an augmented homeostatic T and B cell response to the colonizing bacteria, in that this process does not involve intestinal inflammation. Interestingly, the magnitude of hinge-specific serum IgG is higher in Sweden/US infants than that of Ugandan infants. This might be due to how soon infants raised in different lifestyles are introduced to solid food, and how long breast feeding is continued. However, whether the generation of this adaptive IgG response is simply a result of antigen encounter, or the flagellin-specific antibodies are actively involved in *Lachnospiraceae* colonization needs further investigation. Interestingly, this heightened sero-reactivity to *Lachnospiraceae* flagellins declines with age, in that the flagellin epitope response is hardly detected in healthy adults, consistent with an active regulation of this B cell reactivity and the associated T cell reactivity. It is unclear which immune pathways participate in the downregulation of such immune responses, and the robust IgG sero-reactivity to *Lachnospiraceae* flagellins in CD patients indicates a loss of immunological control. Because flagellins are potent activators of several innate immune receptors, it is possible that defective innate immune pathways are involved. Interestingly, sero-reactivity to the *Lachnospiraceae* flagellin hinge peptide was also found elevated in a subset of patients with severe chronic fatigue syndrome(*64*). These data are consistent with a homeostatic immune response to flagellin in infancy being converted to a pathologic response later in life.

Data in this study raise many interesting and important questions. The strong serum IgG response observed in both healthy infants and CD patients makes us ask what role the hinge epitope specific B cells play in homeostasis and in the early phases of CD. Does the flagellin specific B cell (and CD4^+^ T cell) response in Crohn’s patients decline with age as in most healthy children, and then increase at the onset of the pathology, or does it remain elevated from infancy? Why do flagellin specific IgG antibodies predominantly target the highly conserved hinge peptide and are these antibodies germline or somatically hypermutated? Although we mainly focused on characterizing the flagellin specific sero-reactivity in this study, we are also interested to see whether flagellin specific, especially hinge epitope specific IgG and secretory IgA exist in the intestinal mucosa, and if so, do they block the uptake of flagellins into lymphoid tissues and thus prevent systemic activation? Furthermore, we previously showed that the Treg to T effector ratio of flagellin reactive CD4^+^ cells were significantly reduced in CD patients compared to HCs. This finding leads us to ask whether *Lachnospiraceae* bacteria express flagellins and produce short chain fatty acids such as butyrate synergistically to stimulate predominately Treg cells in most humans, but not in patients with CD. The answers to these questions will await further studies which will be facilitated by the development of the flagellin peptide cytometric bead array that can readily identify patients early in their disease course. Considering that an augmented flagellin-specific CD4^+^ T cell response accompanies multi-flagellin IgG reactivity, identification of CD patients with heightened hinge reactivity might also benefit patient treatment by offering a novel flagellin peptide-guided immunotherapy, which ablates circulating flagellin-specific memory CD4^+^ T cells by concomitant cell activation and metabolism inhibition(*27*).

## MATERIALS AND METHODS

### Study Design

The objective of this research was to identify immunodominant B cell peptide epitopes in the *Lachnospiraceae* bacterial family in patients with Crohn’s disease and in healthy/non-IBD control infants, and correlate this immune reactivity with clinical disease course in the former. This was accomplished by performing a novel flagellin peptide array and flagellin peptide cytometric bead array with an adult discovery cohort containing patients with Crohn’s disease, ulcerative colitis, and healthy controls, and a validation cohort of pediatric treatment-naïve Crohn’s patients and non-IBD controls. Furthermore, a cohort of healthy and non-IBD control infants and their mothers from three distinct geographic locations was utilized for the discovery of the homeostatic antibody response to the same dominant B cell epitope in *Lachnospiraceae* flagellins. Sample sizes were selected on the basis of expected variance and effect size in well-characterized experimental systems and were large enough to achieve a ≥90% power to detect differences >20% among groups. Replicated samples were randomly omitted. Experiments of the validation cohort were conducted blinded until data analyses.

### Patient sample collection

Peripheral blood samples along with medical metadata of IBD patients and healthy volunteers in the UAB cohort were collected at the Kirklin Clinic of UAB Hospital and in research labs in Birmingham, Alabama upon informed consent. All steps followed UAB’s Institutional Review Board guidelines (IRB X300001155, X140515004, and F081003001) and the Declaration of Helsinki Principles. Serum samples of subjects in the RISK cohort were obtained from Dr. Subra Kugathasan at Emory University, USA. Serum samples of subjects in the Uganda and US cohort were obtained from Dr. Edward N. Janoff at the University of Colorado Denver, USA. Serum samples of healthy and allergic subjects in the Sweden cohort were obtained from Dr. Maria C. Jenmalm at Linköping University, Sweden. The Regional Ethics Committee for Human Research at Linköping University approved the study (Dnr 99323). Informed consent was obtained from both parents before inclusion. All serum samples of the RISK, Uganda, and US/Sweden cohorts were obtained through Material Transfer Agreement.

### Motif discovery

Protein sequences (D0-1N p0-75) of *Lachnospiraceae* flagellins included in the flagellin peptide microarray plus *S. dublin* FliC and *E. coli* FliC were analyzed with MEME Suite 5.5.0 for the discovery of recurring protein motifs with fixed length (6-35 width), with 0-order model of sequences used as the background model. Motifs discovered were ordered by E-value and the most significant motif was shown in **Figure 1**.

### Microbiota protein microarray

Microbiota protein microarray was performed as previously described(*24*). In brief, recombinant bacterial flagellin proteins were expressed and diluted in 10mM Tris (PH7.4) with 20% glycerol and 0.1% sodium dodecyl sulfate at 0.2mg/ml. Antigens were printed in quadruplicate on slides coated with nitrocellulose pads (Maine Manufacturing) using a Spotbot Personal microarrayer (ArrayIt). Antigen printed pads were blocked with SuperBlock (ThermoFisher) for 1 hour before probed with serum samples (diluted at 1:100 in SuperBlock). After 1 hour, pads were washed 3 times with PBS plus 0.05% Tween 20, and then incubated with Dylight 550 Goat anti-human IgA (ImmunoReagents, Cat # GtxHu-001-E550NHSX) and Dylight 650 Donkey anti-human IgG (Invitrogen, Cat # SA5-10129) at 1:1000 dilution for 1 hour. Finally, pads were washed 3 times with PBS plus Tween, air-dried, and scanned using GenePix 4000B imager (Axion).

### Flagellin peptide microarray

1512 overlapping 15mer peptides derived from 19 *Lachnospiraceae* flagellins of human and mouse origin were synthesized and printed on individual glass slide as previously described(*65*). Slides were blocked with SuperBlock for 1 hour before they were probed with human sera (dilution factor indicated in corresponding Figure Legends). Slides were washed 3 times with PBS plus 0.05% Tween 20 after 1 hour incubation with serum samples, and then incubated with Dylight 550 Goat anti-human IgA and Dylight 650 Donkey anti-human IgG at 1:1000 dilution for 1 hour. Finally, slides were washed 3 times with PBS plus Tween, air-dried, and scanned using GenePix 4000B imager.

### Generation of flagellin peptide microarray heatmaps

UAB cohort: 1477 of the original 1512 peptides are depicted (columns in the heatmap). A peptide was excluded if more than 190 subjects had signal greater than 2.16 (i.e., log_10_(signal + 1) = .5). 265 of the original 288 subjects are depicted (rows in the heatmap). A subject was excluded if more than 500 peptides had signal greater than 2.16. These thresholds were determined by identifying cutoffs in long tails of distributions subjects and peptides. RISK cohort: 474 peptides and 120 subjects are depicted.

### Flagellin peptide cytometric bead array

8 conserved peptides from *Lachnospiraceae* flagellins were synthesized, with biotin and a GSGSG linker attached to their N-terminus. D1N p41-59 and D0-1N p25-59 include a 50/50 mixture of 2 consensus peptides with one amino acid different, respectively. Sequence of each peptide is listed as follows:

D0N p1-19: [Biotin]GSGSGMVVQHNLRAMNSNRMLGIT[COOH]

D0N p25-44: [Biotin]GSGSGKSTEKLSSGYKINRAADDAA[COOH]

D1N p41-59: [Biotin]GSGSGDDAAGLTISEKMR(S/K)QIRGL[COOH] D0-1N p25-59:

[Biotin]GSGSGKSTEKLSSGYKINRAADDAAGLTISEKMR(S/K)QIRGL[COOH]

D1N p74-102: [Biotin]GSGSGQTAEGALTEVHDMLQRMNELAVQAANGTN[COOH]

D1C p391-407: [Biotin]GSGSGLGAVQNRLEHTINNLDN[COOH]

D0-1C p410-435: [Biotin]GSGSGENTTAAESQIRDTDMATEMVKYSNNN[COOH]

D0C p437-460: [Biotin]GSGSGLAQAGQSMLAQSNQANQGVLSLLG[COOH]

Note that () indicates that it’s a 50/50 mix of peptides at this position.

Flagellin peptide cytometric bead array was performed as previously described(*66*). In brief, biotinylated flagellin peptides were conjugated on to Streptavidin coated 4um polystyrene beads (Spherotech, Inc. Cat # PAK-4067-8K). Beads conjugated with each individual peptide have different fluorescent intensity in the APC/Cy7 channel so that they can be differentiated from each other by flow cytometry. After incubation of 40ul diluted serum samples (serum samples were diluted at 1:200 and 1:600 in SuperBlock to ensure sufficient detection of epitope-specific IgG response; IgM and IgA responses were detected at 1:200 and 1:600 with IgG, respectively) from individuals with CD, UC, and HCs with the array of peptide-bound beads, antibody response to individual peptides was detected with different fluorescently labeled secondary antibodies (Goat anti-human IgM-AF488, SouthernBiotech, Cat # 2020-30; Goat anti-human IgG-PE, SouthernBiotech, Cat # 2040-09; Goat anti-human IgA-PE, SouthernBiotech, Cat # 2050-09; and Goat anti-human IgG-AF488, SouthernBiotech, Cat # 2040-30) using CytoFLEX cytometer (The Beckman Coulter Life Sciences).

### Calculation of epitope-specific antibody concentrations in flagellin peptide cytometric bead array

Biotinylated Goat F(ab)_2_ anti-human isotype antibodies (anti-human IgM, SouthernBiotech, Cat # 2022-01; anti-human IgA, SouthernBiotech, Cat # 2052-01, and anti-human IgG, SouthernBiotech, Cat # 2042-01) were conjugated on to selected peaks of Streptavidin coated 4um polystyrene capture beads, respectively, and incubated with 0.75x serial dilutions of purified polyclonal human antibodies (IgG, SouthernBiotech, Cat # 0150-01; IgM, SouthernBiotech, Cat # 0158L-01; IgA, SouthernBiotech, Cat # 0155K-01) at known concentrations. Standard curves were obtained based on MFI detected by secondary antibodies bound using CytoFLEX cytometer. Flow cytometry standard (FCS) files of both samples and the standard curve were applied to a set of MATLAB programs (https://github.com/UAB-Immunology-Institute/cba-toolbox) generated by Dr. Rosenberg at UAB for the quantification of peptide-specific antibodies in the corresponding isotype(*66*).

### Statistical analysis

Statistical analyses were performed with GraphPad PRISM, version 9.5, with nonparametric Kruskal-Wallis test or two-way ANOVA as indicated. Dunn’s or Tukey’s multiple comparisons test was used when comparing multiple groups. All statistical tests were 2-sided. Specific tests used and values of n are stated in corresponding figure legends. Data were presented as means ± SEM. The differences were considered statistically significant at P<0.05 (*P<0.05, **P<0.01, ***P<0.001, ****P<.0001) and ns indicates not significant.

## Acknowledgments

We thank all the patients and volunteers that participated in this study and the hospital staff for obtaining blood samples from participants. We thank all site investigators from 28 sites and the Crohn’s and Colitis Foundation of America for supporting the RISK cohort study. The authors would also like to thank BioRender for providing the platform for figure generation.

## Authorship Contributions

COE initiated, designed, and supervised the study. QZ designed and conducted the study, analyzed the data, and wrote the manuscript. LWD participated in study design, conducted the microbiota protein microarray and flagellin peptide microarray, and analyzed the data. JTK and RGK participated in the setup and data analysis of the flagellin peptide cytometric bead array. AFR analyzed and visualized the data, and designed the program for the quantification of epitope-specific antibodies in the cytometric bead array. PJM, LAD, ENJ, MCJ and COE acquired blood samples and clinical diagnoses from study participants.

