## Supplementary Figures for "Crohn’s patients and healthy infants share immunodominant B cell response to commensal flagellin peptide epitopes"

**Zhao *et al*. Supplementary Figure 1**

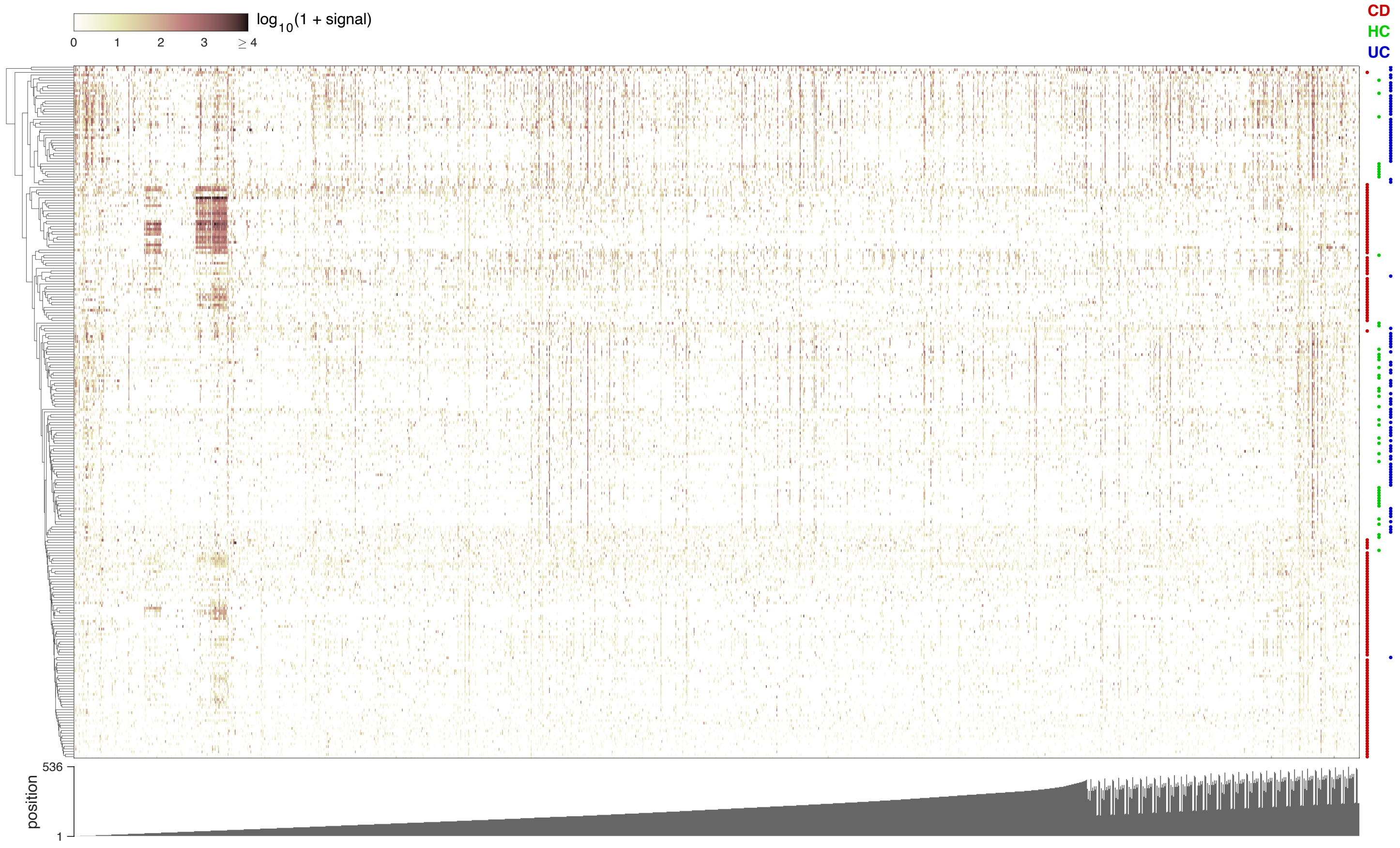


**Supplementary Figure 1.** Heatmap showing serum IgG reactivity for all fifteen-residue flagellin peptides (including 108 positions derived from 19 different flagellins) in CD, HC, and UC subjects. Method for clustering and sorting are the same as in **Figure 1B**. Cohort groups are indicated by colored dots in the right margin. Peptide starting position (relative to the N-terminus) is indicated on the bottom (peptides in the C-terminus were sorted based on their sequential alignment from the end of the C-terminus).

**Zhao *et al*. Supplementary Figure 2**

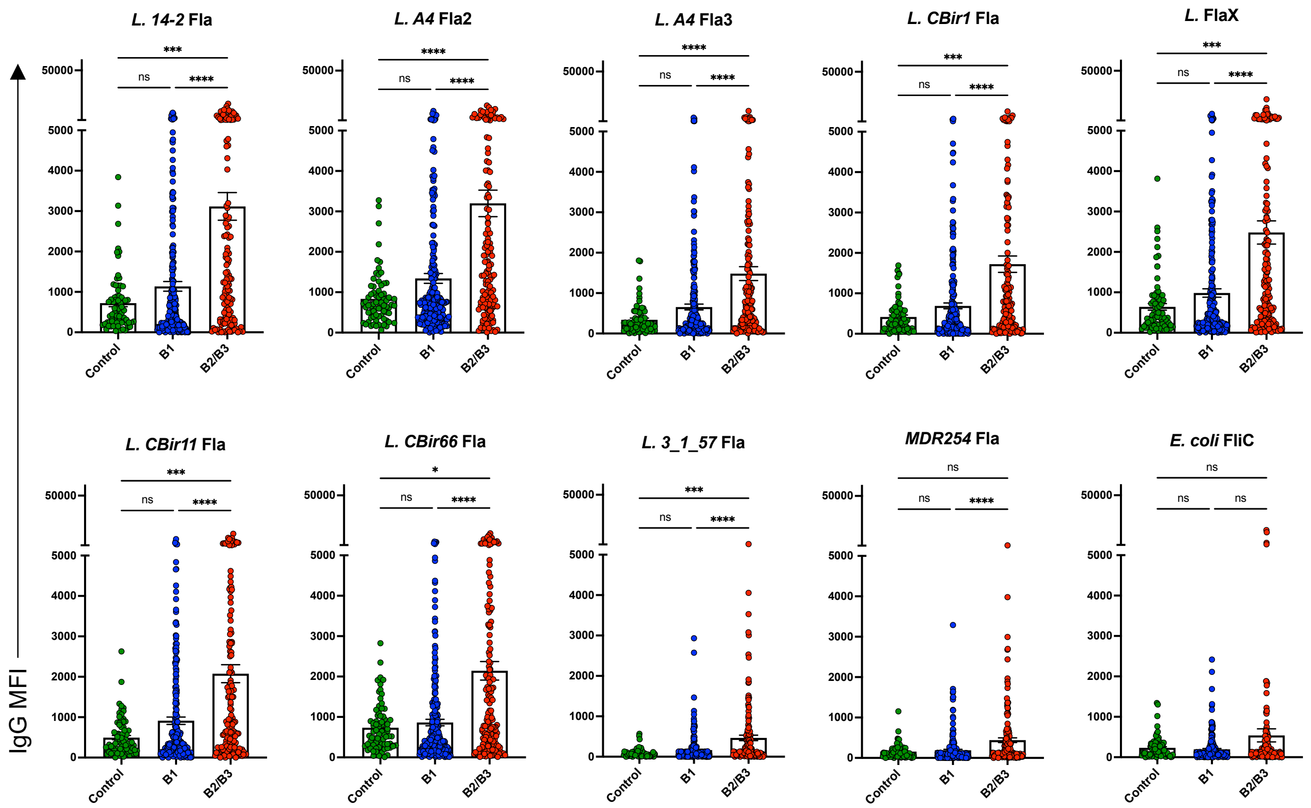


**Supplementary Figure 2.** Sera of CD patients (grouped by disease behavior at 3-year follow-up, n=250 for B1, n=142 for B2/3) and non-IBD controls (n=72) from the RISK cohort were probed against the microbiota protein microarray. MFI of serum IgG specific for indicated flagellin antigens are presented as means ± SEM and analyzed with nonparametric Kruskal-Wallis test. *P<0.05, **P<0.01, ***P<0.001, ****P<0.0001.

**Zhao *et al*. Supplementary Figure 3**

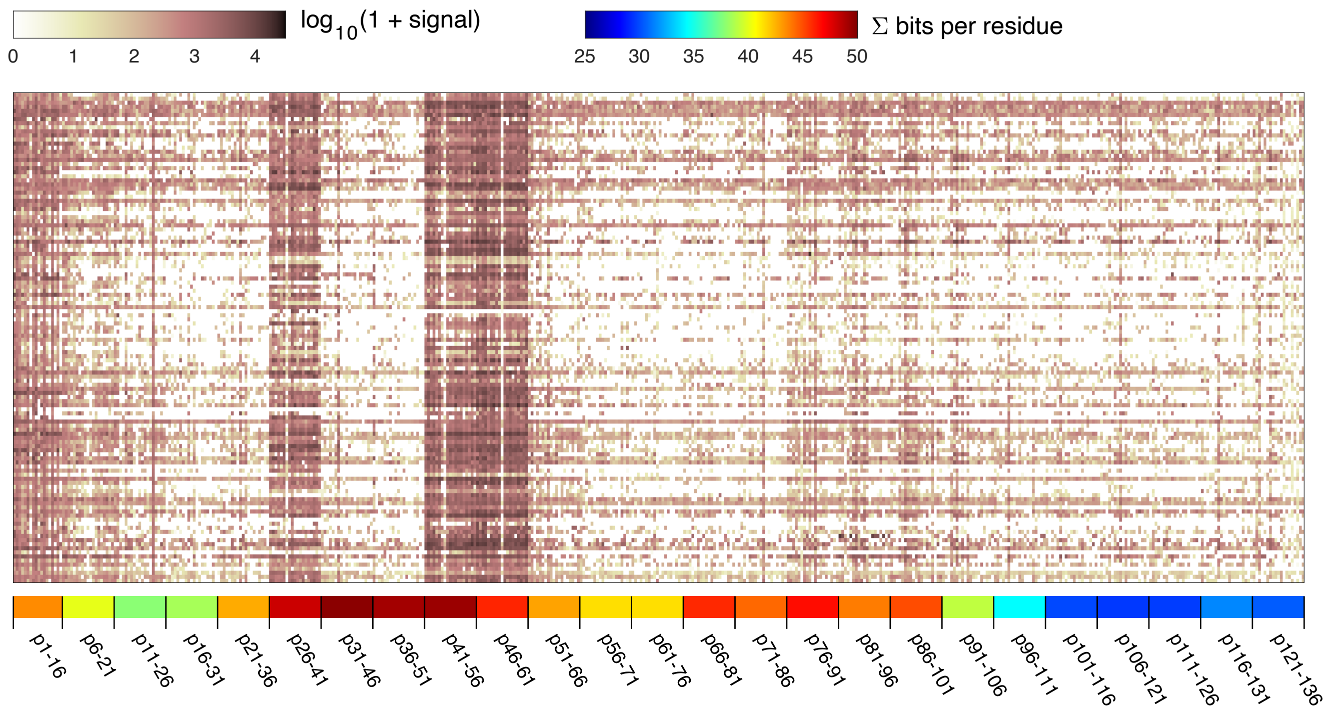


**Supplementary Figure 3.** Heatmap showing peptide reactivity data for fifteen-residue flagellin peptides (including 25 positions and 19 different flagellins) for multi-flagellin reactive CD subjects in the RISK cohort. Sera were diluted at 1:30. Each row represents reactivity of an individual subject (not clustered), whereas columns represent peptides sorted by peptide position (and include up to 19 flagellins per position), and color indicates log-transformed signal. Peptide starting position (relative to the N-terminus) is indicated on the bottom. For each 15-mer segment, per-residue information content is computed at each position across all flagellins (except *L. CBir11* Fla and *L. CBir66* Fla) and summated to indicate sequence conservation, shown as color.

**Zhao *et al*. Supplementary Figure 4**

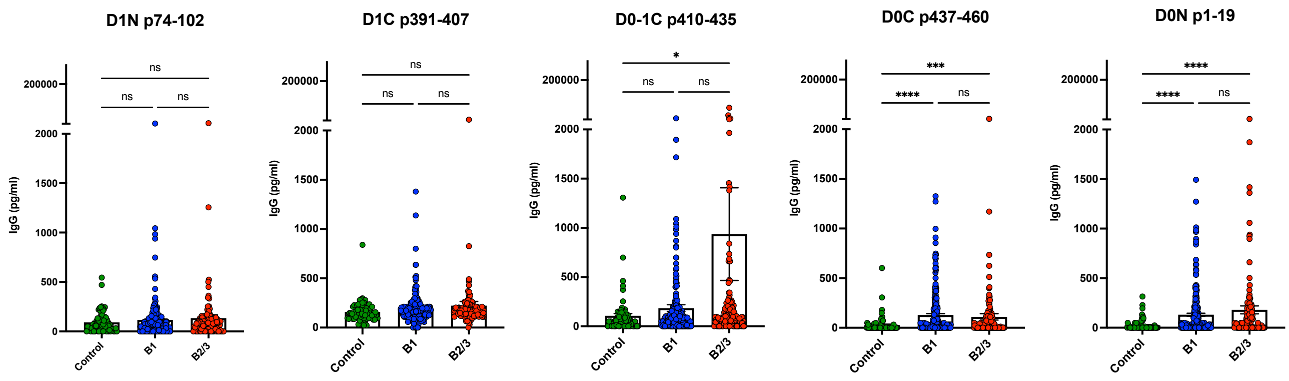


**Supplementary Figure 4.** Sera of CD patients (grouped by disease behavior at 3-year follow-up, n=249 for B1, n=140 for B2/3) and non-IBD controls (n=72) from the RISK cohort were probed against the flagellin peptide cytometric bead array. Epitope-specific IgG concentration in different groups to indicated flagellin peptide epitopes is shown. Data are presented as means ± SEM and analyzed with nonparametric Kruskal-Wallis test. *P<0.05, **P<0.01, ***P<0.001, ****P<0.0001.

**Zhao *et al*. Supplementary Figure 5**

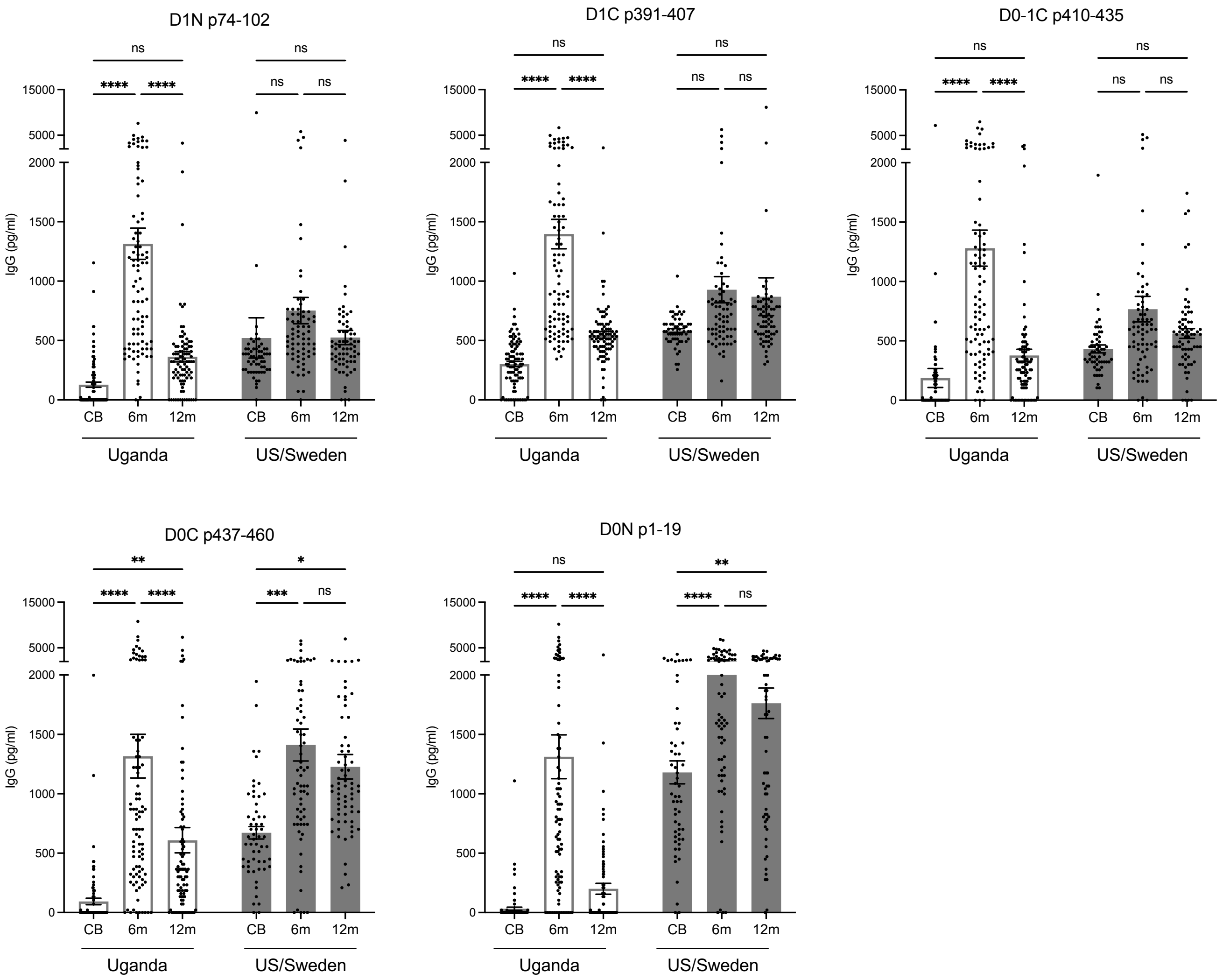


**Supplementary Figure 5.** Sera of healthy and non-IBD control infants at indicated timepoints from the Uganda cohort (n=92) and the US (n=16) /Sweden (n=54) cohort were probed against the flagellin peptide cytometric bead array. IgG concentration specific to indicated *Lachnospiraceae* peptide epitopes in different groups is shown. Data are presented as means ± SEM and analyzed with two-way ANOVA and Tukey's multiple comparisons test. *P<0.05, **P<0.01, ***P<0.001, ****P<0.0001.
